# Evidence for reassortment of highly divergent novel rotaviruses from bats in Cameroon, without evidence for human interspecies transmissions

**DOI:** 10.1101/054072

**Authors:** Claude Kwe Yinda, Mark Zeller, Nádia Conceição-Neto, Piet Maes, Ward Deboutte, Leen Beller, Elisabeth Heylen, Stephen Mbigha Ghogomu, Marc Van Ranst, Jelle Matthijnssens

## Abstract

Bats are an important reservoir for pathogenic human respiratory and hemorrhagic viruses but only little is known about bat viruses causing gastroenteritis in humans, including rotavirus A strains (RVA). Only three RVA strains have been reported in bats in Kenya (straw-colored fruit bat) and in China (lesser horseshoe and a stoliczka’s trident bat), being highly divergent from each other. To further elucidate the potential of bat RVAs to cause gastroenteritis in humans we started by investigating the genetic diversity of RVAs in fecal samples from 87 straw-colored fruit bats living in close contact with humans in Cameroon using metagenomics. Five samples contained significant numbers of RVA Illumina reads, sufficient to obtain their (near) complete genomes. A single RVA strain showed a close phylogenetic relationship with the Kenyan bat RVA strain in six gene segments, including VP7 (G25), whereas the other gene segments represented novel genotypes as ratified by the RCWG. The 4 other RVA strains were highly divergent from known strains (but very similar among each other) possessing all novel genotypes. Only the VP7 and VP4 genes showed a significant variability representing multiple novel G and P genotypes, indicating the frequent occurrence of reassortment events.

Comparing these bat RVA strains with currently used human RVA screening primers indicated that several of the novel VP7 and VP4 segments would not be detected in routine epidemiological screening studies. Therefore, novel VP6 based screening primers matching both human and bat RVAs were developed and used to screen samples from 25 infants with gastroenteritis living in close proximity with the studied bat population. Although RVA infections were identified in 36% of the infants, Sanger sequencing did not indicate evidence of interspecies transmissions.

This study identified multiple novel bat RVA strains, but further epidemiological studies in humans will have to assess if these viruses have the potential to cause gastroenteritis in humans.

## Introduction

Rotaviruses (RVs) are major enteric pathogens causing severe dehydrating diarrhea mostly in juvenile humans and animals worldwide (1). RVs belong to the family *Reoviridae* and the genus *Rotavirus* consists of eight species (A-H) (2). Recently, a distinct canine RV species has been identified and tentatively named *Rotavirus I* (RVI), although ratification by the International Committee on Taxonomy of Viruses (ICTV) is pending (3). Group A rotaviruses (RVAs) are the most common of all rotavirus species and infect a wide range of animals including humans (4). The rotavirus genome consists of 11 double-stranded RNA segments encoding 6 structural viral proteins (VP1–VP4, VP6, and VP7) and 6 nonstructural proteins (NSP1–NSP6). In 2008, a uniform classification system was proposed for each of the 11 gene segments resulting in G-, P-, I-, R-, C-, M-, A-, N-, T-, E-and H-genotypes for the VP7, VP4, VP6, VP1-VP3, NSP1-NSP5 encoding gene segments, respectively (5, 6). This classification system enabled researchers to compare genotype constellations between different host species and infer evolutionary patterns. In recent years host specific genotype constellations have been determined for pigs, ruminants, horses and cats/dogs (7–10).

A rich but, until recently, underappreciated reservoir of emergent viruses are bats. They make up to 20% of the ~5,500 known terrestrial species of mammals (11) and are the second most abundant mammals after rodents (12). Several viruses pathogenic to humans are believed to have originated from bats, including Severe Acute Respiratory Syndrome (SARS), Middle East Respiratory Syndrome (MERS)-related coronaviruses, as well as *Filoviridae*, such as Marburgvirus, and Henipaviruses, such as Nipah and Hendra virus (13–15). In the last decades advances in viral metagenomics including high-throughput next-generation sequencing technologies have led to the discovery of many novel viruses including enteric viruses from bats (15, 16). However, rotaviruses have only been reported sporadically in bats and only three RVA strains have been characterized so far. The first strain was reported in a straw-colored fruit bat (*Eidolon helvum)* in Kenya. This partially sequenced strain was named RVA/Bat-wt/KEN/KE4852/2007/G25P[6], and possesses the following genotype constellation: G25-P[6]-I15-Rx-C8-Mx-Ax-N8-T11-E2-H10 (17). Two other bat RVAs were found in China in a lesser horseshoe bat (*Rhinolophus hipposideros*) and a stoliczka’s trident bat (*Aselliscus stoliczkanus*) named RVA/Bat-tc/CHN/MSLH14/2012/G3P[3] and RVA/Bat-tc/CHN/MYAS33/2013/G3P[10], respectively (18, 19). Phylogenetic analysis showed that strains MSLH14 and MYAS33, although sampling sites were more than 400 km apart, shared the same genotype constellation (G3-P[x]-I8-R3-C3-M3-A9-N3-T3-E3-H6) except for the P genotype which was P[3] for MSLH14 and P[10] for MYAS33.

To further study the genomics of RVA in bats and their zoonotic potential in humans, we screened stool samples of straw-colored fruit bats (*Eidolon helvum*) living in close proximity with humans in the South West Region of Cameroon, as well as samples from infants with gastroenteritis. Our choice of this region is due to the fact that bats are considered a delicacy and the species sampled are the most commonly eaten bat species in these localities.

## Methods

### Bat sample collection

Bat samples were collected between December 2013 and May 2014 using a previously described method (20), after obtaining administrative authorization from the Delegation of Public Health for South West Region. Briefly, bats were captured in 3 different regions (Lysoka, Muyuka and Limbe) of the South West Region of Cameroon (Fig. 1) using mist nets around fruit trees and around human dwellings. Captured bats were retrieved from the traps and held in paper sacks for 10-15 min, allowing enough time for the excretion of fresh fecal boluses. Sterile disposable spatulas were used to retrieve feces from the paper sacks, and placed into tubes containing 1 ml of universal transport medium (UTM, Copan Diagnostics, Brescia, Italy). Labeled samples were put on ice and then transferred to the Molecular and cell biology laboratory, Biotechnology Unit, University of Buea, Cameroon and stored at −20°C, until they were shipped to the Laboratory of Viral Metagenomics, Leuven, Belgium where they were stored at −80°C. Each captured bat was assessed to determine species, weight (g), forearm length (mm), sex, reproductive state, and age. All captured bats were then marked by hair clipping to facilitate identification of recaptures, and released afterwards. Trained zoologists used morphological characteristics to determine the species of the bats before they were released. No clinical signs of disease were noticed in any of these bats.

**Figure 1:**
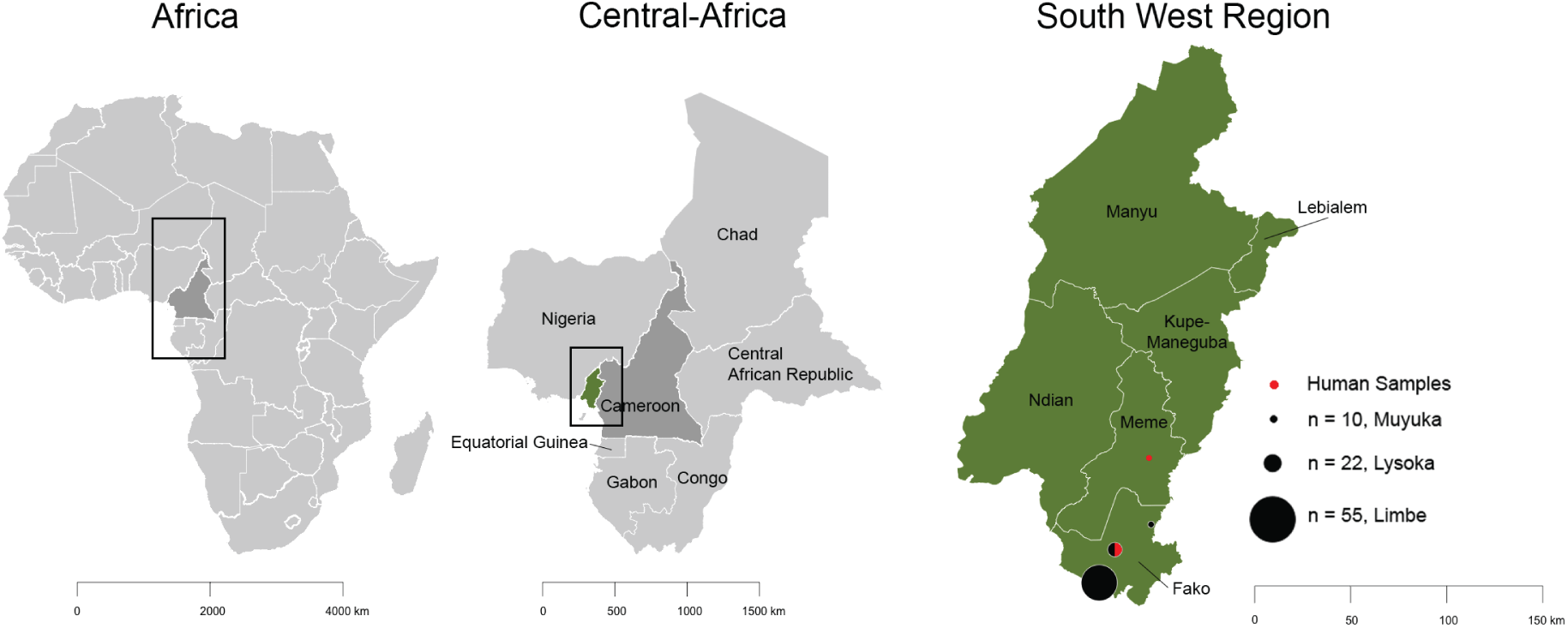
Map of study site (South West Region, Cameroon). The number of bat (black filled circles) and human (red filled circles,) samples are indicated.

### Sample preparation for NGS

Eighty-seven fecal samples were grouped into 24 pools each containing three to five samples and treated to enrich viral particles as follows: fecal suspensions were homogenized for 1 min at 3000 rpm with a MINILYS homogenizer (Bertin Technologies, Montigny-le-Bretonneux, France) and filtered consecutively through 100 μm, 10 μm and 0.8 μm membrane filters (Merck Millipore, Massachusetts, USA) for 30 s at 1250 *g*. The filtrate was then treated with a cocktail of Benzonase (Novagen, Madison, USA) and Micrococcal Nuclease (New England Biolabs, Massachusetts, USA) at 37°C for 2h to digest free-floating nucleic acids. Nucleic acids were extracted using the QIAamp Viral RNA Mini Kit (Qiagen, Hilden, Germany) according to the manufacturer’s instructions but without addition of carrier RNA to the lysis buffer. First and second strand cDNA synthesis was performed and random PCR amplification for 17 cycles were performed using a Whole Transcriptome Amplification (WTA) Kit procedure (Sigma-Aldrich), with a denaturation temperature of 95°C instead of 72°C to allow the denaturation of dsDNA and dsRNA. WTA products were purified with MSB Spin PCRapace spin columns (Stratec, Berlin, Germany) and the libraries were prepared for Illumina sequencing using the NexteraXT Library Preparation Kit (Illumina, San Diego, USA). A cleanup after library synthesis was performed using a 1.8 ratio of Agencourt AMPure XP beads (Beckman Coulter, Inc., Nyon, Switzerland) (21). Sequencing of the samples was performed on a HiSeq 2500 platform (Illumina) for 300 cycles (2x150 bp paired ends). Partial sequences were completed using RT-PCRs with specific primers (Supplementary table S1). For gene segments lacking the 5′ and/or 3′ ends of the ORF the single primer amplification method (Primers in Supplementary table, S1) was used as described previously (22). Sanger sequencing was done on an ABI Prism 3130 Genetic Analyzer (Applied Biosystems, Massachusetts, USA).

### Human fecal sample collection and Screening

Human fecal samples were collected from Lysoka local clinic and Kumba District Hospital of the South west Region of Cameroon (Fig. 1) from patients who were either diarrheic or came into contact with bats directly (by eating, hunting or handling) or indirectly (if family member is directly exposed to bats). The samples were put in UTM containing tubes and stored the same way like the bat samples. Screening primers (suppl. S1) were designed from a consensus sequences of human and bat VP6 RVAs and a total of 25 samples from infants (0-3 years) who had diarrhea were screened by reverse transcriptase polymerase chain reaction (RT-PCR) using the OneStep RT-PCR kit (Qiagen). The products of positive samples were sequenced using Sanger sequencing.

### Genomic and phylogenetic analysis

Raw Illumina reads were trimmed for quality and adapters using Trimmomatic (23), and were *de novo* assembled into Scaffold using SPAdes (24). Scaffolds were classified using DIAMOND in sensitive mode (25). Contigs assigned to RVA were used to map the trimmed reads using the Burrows-Wheeler Alignment tool (BWA) (26). Open reading frames (ORF) were identified with ORF Finder analysis tool (http://www.ncbi.nlm.nih.gov/gorf/Orfig.cgi) and the conserved motifs in the amino acid sequences were identified with HMMER (27). Amino acid alignments of the viral sequences and maximum likelihood phylogenetic trees were constructed in MEGA6.06, (28) using the GTR+G (VP1, VP6, NSP2 and NSP3), GTR+G+I (VP2-VP4, VP7 and NSP1), HKY (NSP4) and T92 (NSP5) substitution models (after testing for the best DNA/protein model), with 500 bootstrap replicates. Nucleotide similarities were also computed in MEGA by pairwise distance using p-distance model. Sequences used in the phylogenetic analysis were representatives of each genotype and the sequences of the RVA discovered in this study. All sequences from the novel viruses were submitted to GenBank.

## Results

### Sample characterization

A total of 24 pools of 3-5 bat fecal samples were constituted, enriched for viral particles and sequenced. Illumina sequencing yielded between 1.1 and 7.8 million reads per pool, and DIAMOND classification (25) of the obtained contigs indicated that five pools contained a significant amount of RVA sequence reads. The percentage reads mapping to RVA in each pool ranged from 0.1-2.4% (Table 1). Partial segments were completed by regular PCR and Sanger sequencing, to obtain at least the entire ORF for each of the obtained variants. Obtained sequences were used for phylogenetic comparison with a selection of representative members of each genotype. The RVA strains discovered in this study were named RVA/Bat-wt/CMR/BatLi08/2014/G31P[42], RVA/Bat-wt/CMR/BatLi09/2014/G30P[42], RVA/Bat-wt/CMR/BatLi10/2014/G30P[42], RVA/bat-wt/CMR/BatLy03/2014/G25P[43] and RVA/Bat-wt/CMR/BatLy17/2014/G30P[xx] hereafter referred to as BatLi08, BatLi09, BatLi10, BatLy03 and BatLy17, respectively. All the obtained sequences were highly divergent from established genotypes and were therefore submitted to the Rotavirus Classification Working group (RCWG) for novel genotype assignments (see below) except for the VP4 gene segment of BatLy17 of which parts of the 3’-end are missing despite repeated attempts to obtain the missing sequence information. However this gene segment is also quite divergent and potentially represent a new genotype.

**Table 1:** Geographic location of sample collection, number of reads per pool, reads mapping to RVAs and reads mapping per gene segment.

| Pool | BatLi10 (P10) | BatLyP03 (P03) | BatLi08 (P08) | BatLi09 (P09) | BatLyP17(P17) |
| --- | --- | --- | --- | --- | --- |
| Location | Limbe | Lysoka | Limbe | Limbe | Lysoka |
| N° of reads/pool | 7,819,081 | 920,217 | 2,140,494 | 1,106,398 | 4,446,078 |
| N° of reads mapping to RVAs | 86,063 | 1,881 | 50,332 | 16,632 | 3,485 |
| Percentage RVA reads | 1.1 | 0.2 | 2.35 | 1.5 | 0.1 |
| N° of reads mapping to VP7 | 1,387 | 145 | 1,775 | 702 | 59 |
| N° of reads mapping to VP4 | 15,513 | 217 | 8,532 | 2,971 | 619 |
| N° of reads mapping to VP6 | 7,520 | 353 | 4,120 | 3,214 | 465 |
| N° of reads mapping to VP1 | 11,667 | 265 | 8,189 | 1,460 | 685 |
| N° of reads mapping to VP2 | 12,727 | 251 | 8,934 | 2,046 | 353 |
| N° of reads mapping to VP3 | 11,552 | 112 | 7,062 | 1,208 | 383 |
| N° of reads mapping to NSP1 | 16,200 | 150 | 5,552 | 2,386 | 554 |
| N° of reads mapping to NSP2 | 2,703 | 68 | 1,924 | 675 | 146 |
| N° of reads mapping to NSP3 | 4,855 | 159 | 2,352 | 1,224 | 100 |
| N° of reads mapping to NSP4 | 957 | 103 | 841 | 390 | 64 |
| N° of reads mapping to NSP5 | 672 | 18 | 341 | 356 | 22 |

### Phylogenetic analysis

The VP7 gene of BatLy03 was 96% identical (on the nucleotide (nt) level) to the Kenyan bat RVA strain KE4852 counterpart, which had been previously classified as a G25 genotype (Fig. 2). BatLi10 and BatLi09 were 99% identical and also clustered closely with strain BatLy17 (92% similar). This cluster was only distantly related to all other known VP7 RVA sequences as well as to strain BatLi08, which also formed a unique long branch in the phylogenetic tree. Both clusters only show similarities below 60% with established genotypes (Fig. 2). The VP7 of these 4 strains (BatLi10, BatLi09, BatLy17 and BatLi08) did not belong to any of the established RVA G-genotypes, according to the established criteria (6), and were assigned genotypes G30 (BatLi09, BatLi10 and BatLy17) and G31 (BatLi08) by the RCWG. For VP4, VP1 and VP3, all five Cameroonian bat RVAs strains were distantly related to other known RVA strains, including the Kenyan and Chinese RVA strains and were therefore assigned to novel genotypes according to the RCWG classification criteria (Fig. 3). The VP4 gene of strains BatLi08, BatLi09 and BatLi10 (representatives of the novel genotype P[42]) were almost 98-100% identical to each other and only 56-75% identical to any other P-genotype. Strains BatLy03 and BatLy17 had 30% nt dissimilarity to each other and their nt identity ranged from 59-76% to other P-genotypes. Only for BatLy03 we were able to obtain the complete VP4 ORF, resulting in its classification as P[43], whereas BatLy17 remained P-unclassified. The VP1 and VP3 genes of BatLi08, BatLi09, BatLi10 and BatLy17 were nearly identical (nt identity range 97-100%) and clustered together but distinct from other established R and M-genotype, thereby representing the new genotypes R15 and M14, respectively (Fig. 3). The VP1 and VP3 genes of BatLy03 were only distantly related to the other four Cameroonian bat RVAs (67-75% nt identity) and are the sole member of the newly assigned genotypes R16 and M15, respectively (Fig. 3). The VP6, VP2, NSP2, NSP3 and NSP5 gene segments of 4 of our strains (BatLi08, BatLi09, BatLi10 and BatLy17) were distantly related to their counterparts of other mammalian and avian RVAs (Fig. 4). For all the 4 strains, these gene segments clustered together and were 98-100% identical to each other and consequently they constitute new genotypes for the different gene segments (I22, C15 N15, T17 and H17, respectively). The VP6, VP2, NSP2, NSP3 and NSP5 gene segments of BatLy03 phylogenetically clustered together with the Kenyan bat RVA strain KE4852 in the previously established I15, C8, N8, T11 and H10 genotypes, respectively (Fig. 4). For NSP1, the Cameroonian bat strains BatLi08, BatLi09, BatLi10 and BatLy17 clustered closely together (98-100% nucleotide sequence identity) in the novel genotype A25, and showed only 67% nucleotide similarity to strain BatLy03 (A26). These 5 new NSP1 gene segments were only 39-40% identical to that of the most closely related established NSP1 genotype A9 (containing the Chinese bats) (Fig. 5). The NSP4 gene segments of all the 5 RVAs discovered in this study were quite divergent to those of other known bat rotaviruses (at most 69% nucleotide sequence identity) and other RVAs (approximately 45-68% nucleotide similarity) forming two distinct clusters. The NSP4 gene segments of strains BatLi08, BatLi09, BatLi10 and BatLy03 (genotype E22) were 98-100% identical but all were 37-38% divergent from that of BatLy03 (E23, Fig. 5).

**Figure 2:**
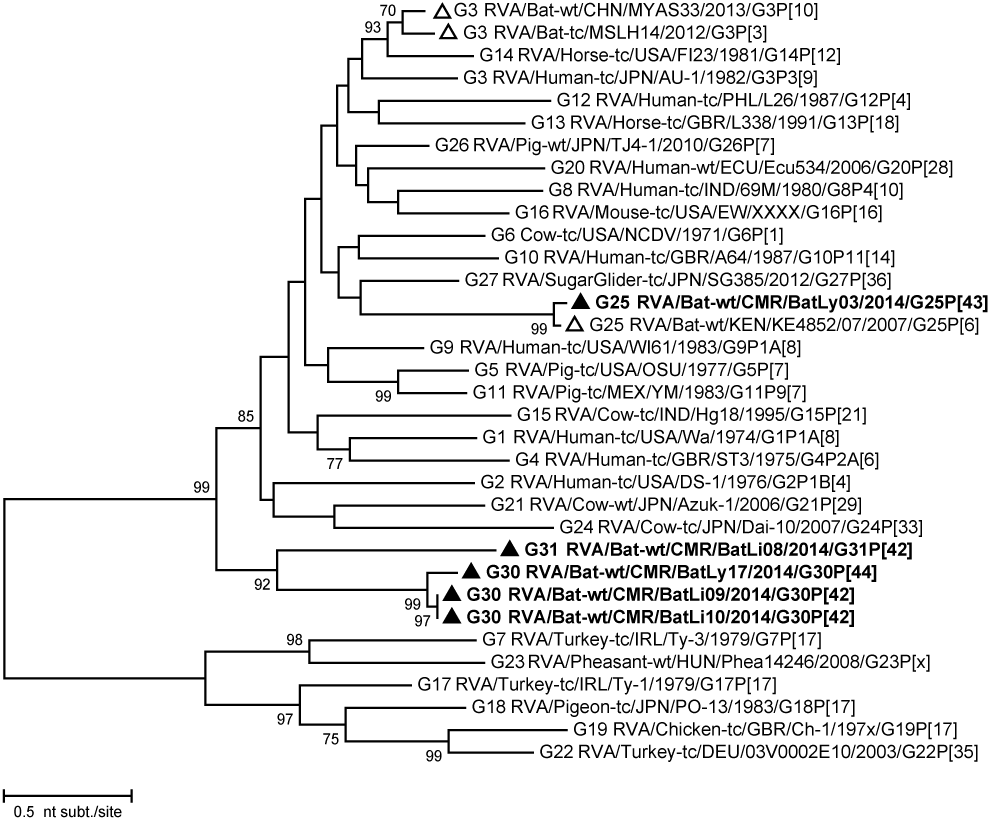
Phylogenetic trees of full-length ORF nucleotide sequences of RVA VP7. Filled triangle: Cameroonian bat RVA strains; open triangles: previously described bat RVA strains. Bootstrap values (500 replicates) above 70 are shown.

**Figure 3:**
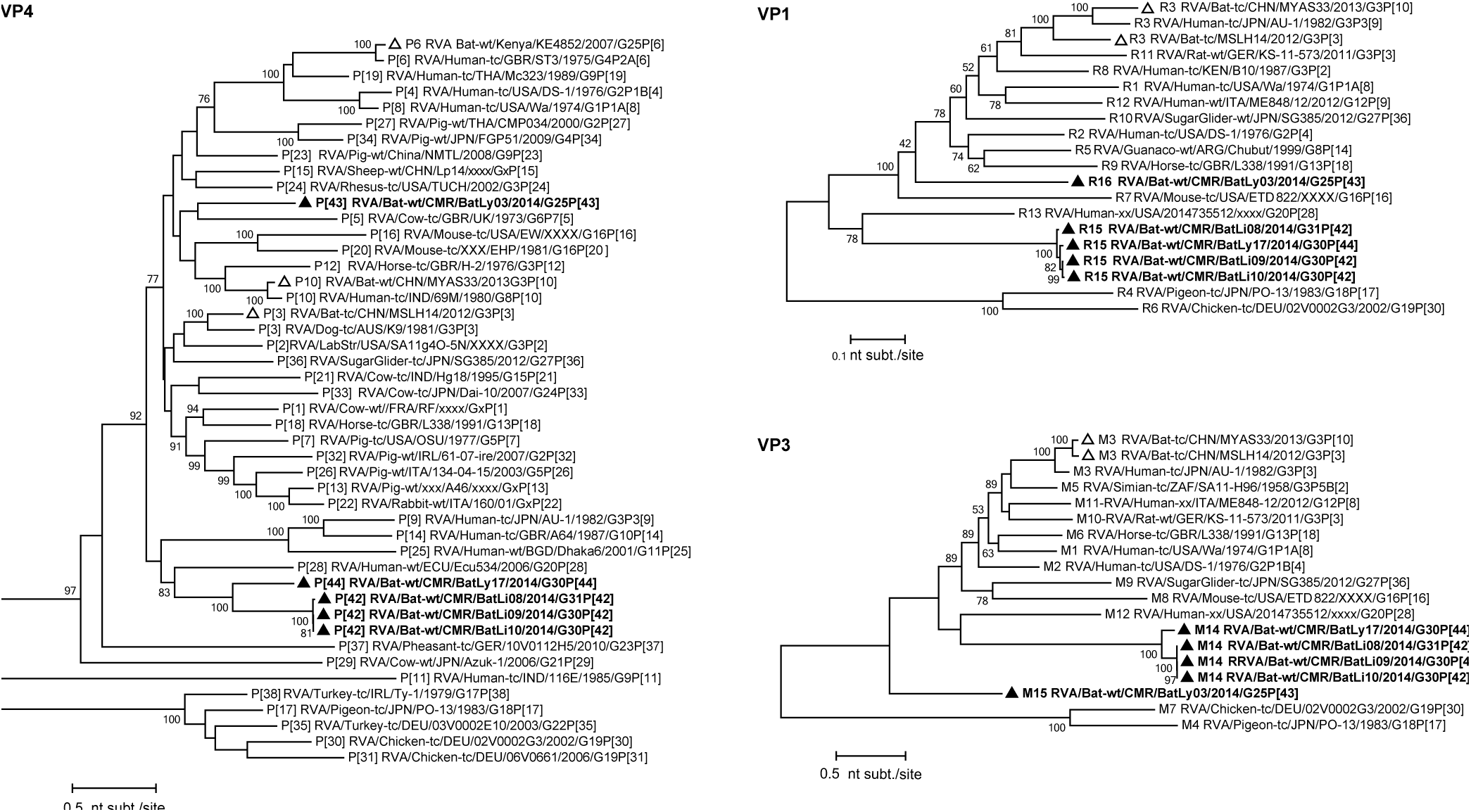
Phylogenetic trees of full-length ORF nucleotide sequences of the RVA VP4, VP1, VP3 gene segments. Filled triangle: Cameroonian strains; open triangles: previously described bat RVA strains. Bootstrap values (500 replicates) above 70 are shown.

**Figure 4:**
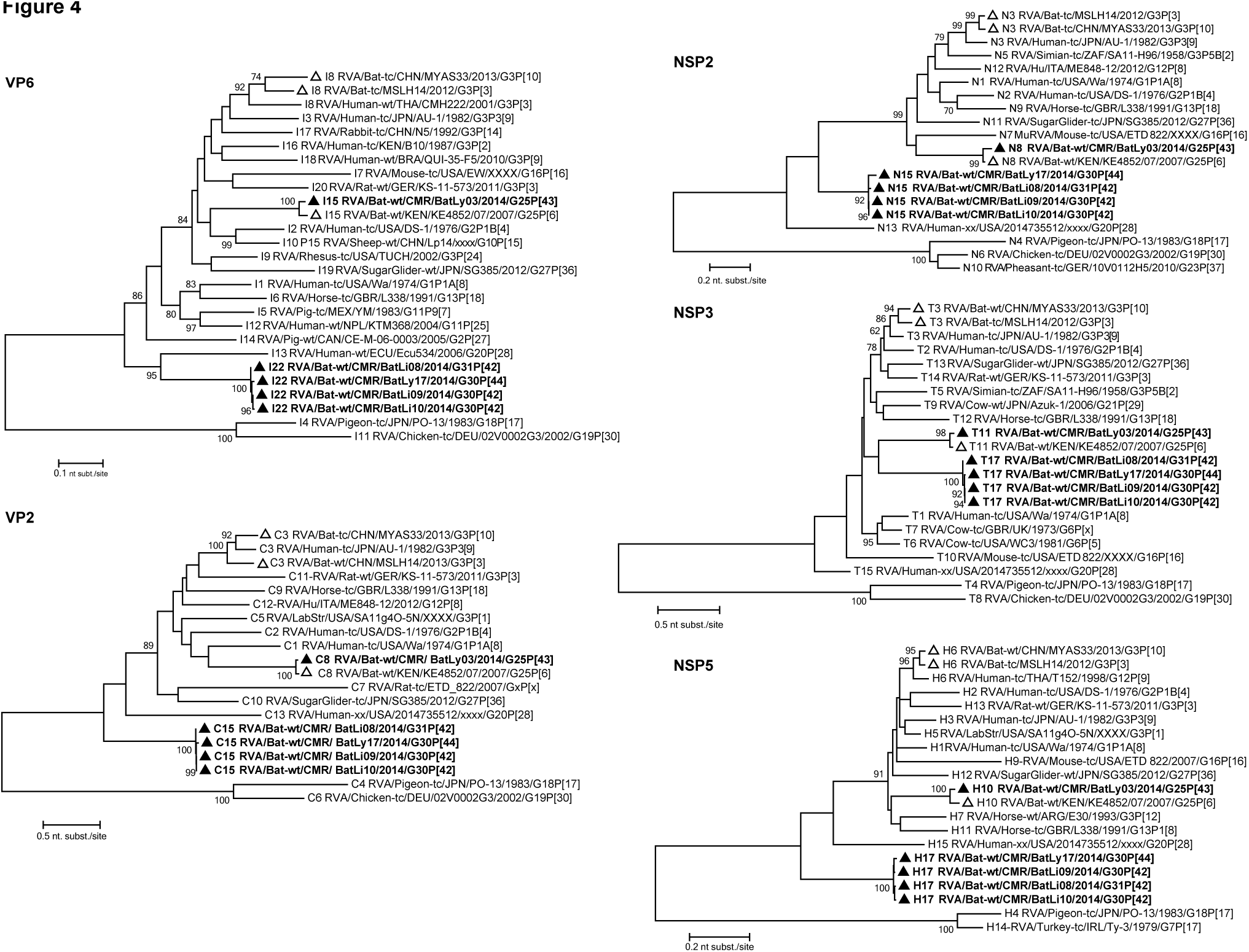
Phylogenetic trees of full-length ORF nucleotide sequences of the RVA VP6, VP2 NSP2, NSP3 and NSP5 gene segments. Filled triangle: Cameroonian strains; open triangles: previously described bat RVA strains. Bootstrap values (500 replicates) above 70 are shown.

**Figure 5:**
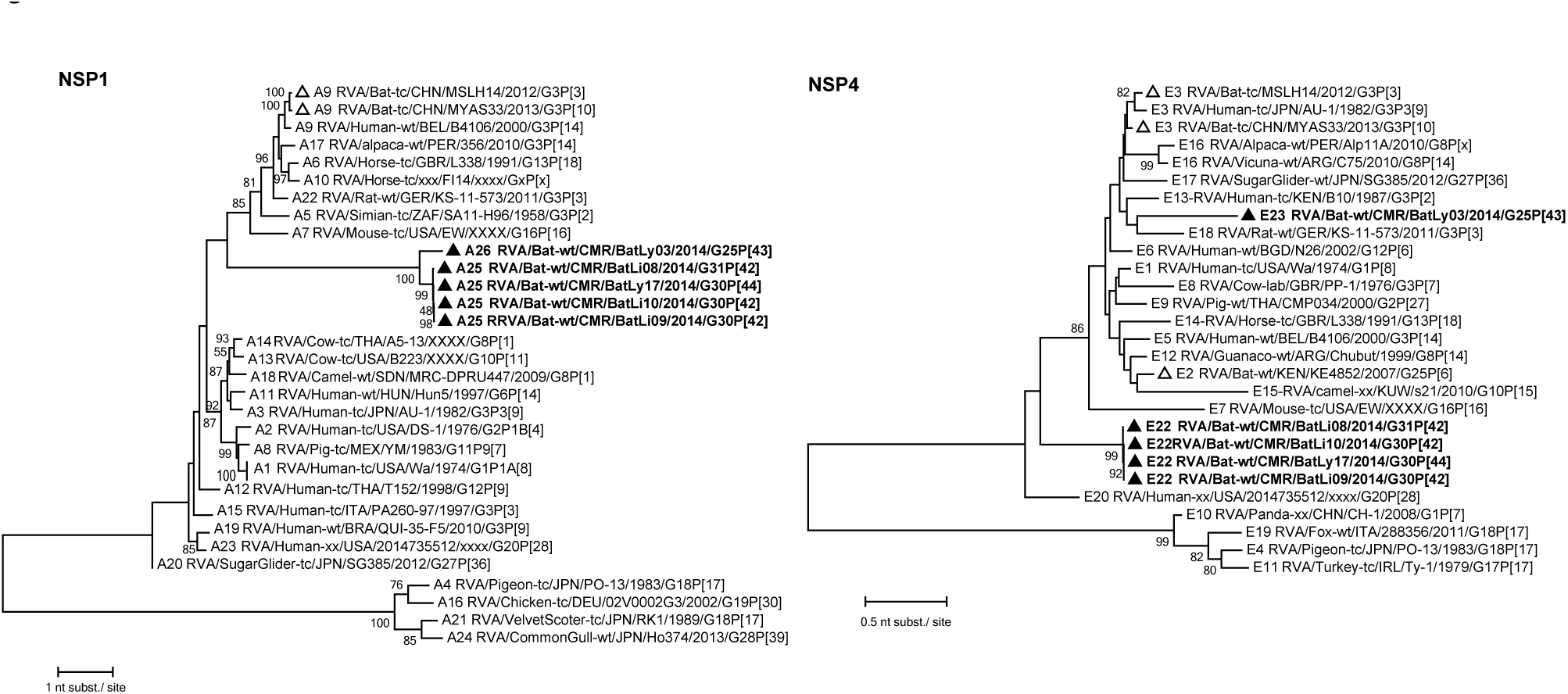
Phylogenetic trees of full-length ORF nucleotide sequences of the RVA NSP1 and NSP4 gene segments. Filled triangle: Cameroonian strains; open triangles: previously described bat RVA strains. Bootstrap values (500 replicates) above 70 are shown.

### Bat rotaviruses in humans?

Several different primer pairs are currently being used to detect human RVA VP7 and VP4 gene segments, to determine the G- and P-genotypes using sequencing or multiplex PCR assays (29–32). In order to find out if the currently used human RVA screening primers would detect the bat RVA strain from this study in case of zoonosis, we compared these primers with their corresponding sequences in the respective gene segments Table 2 (and Suppl. S2).

**Table 2:**
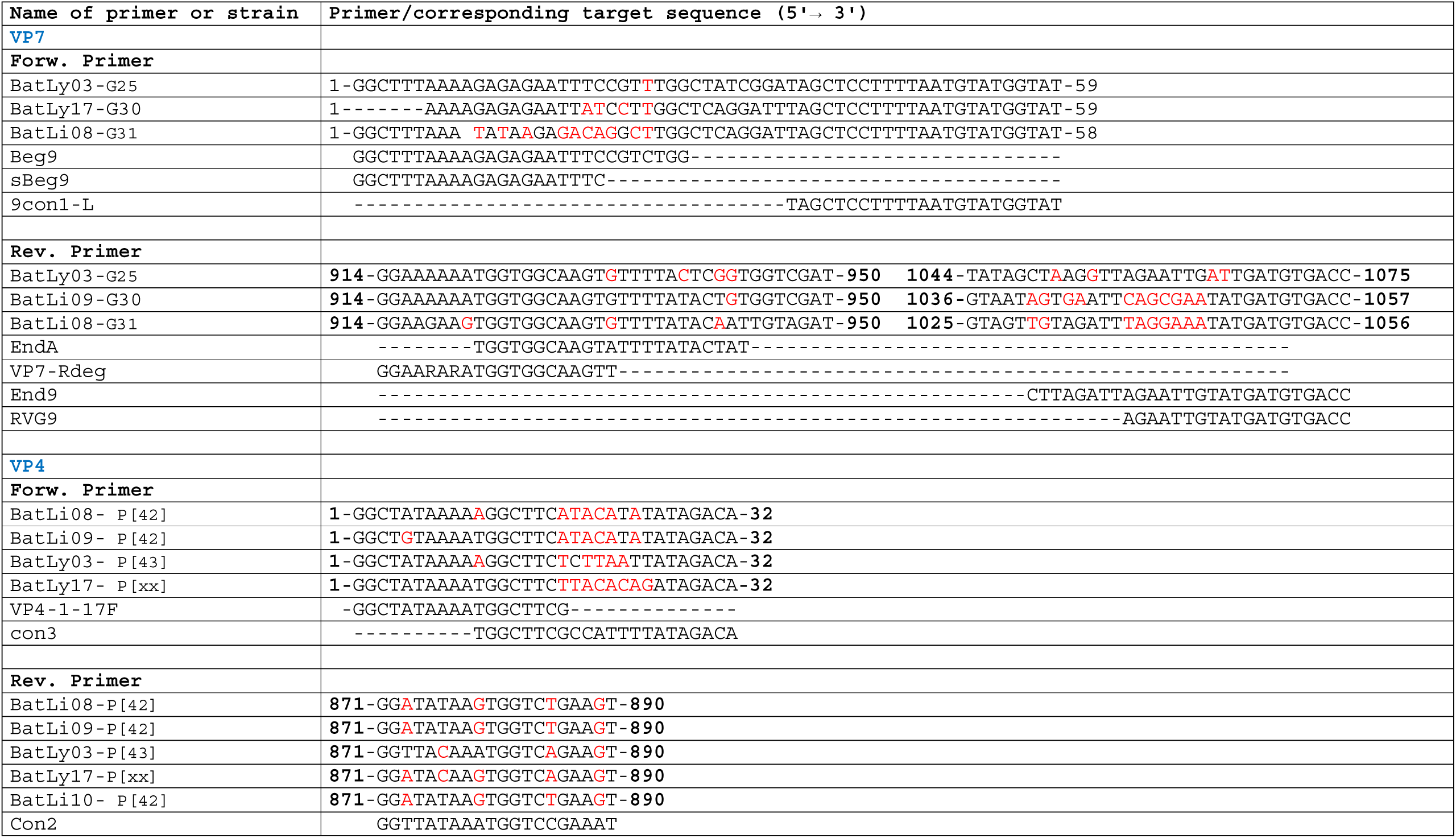
Nucleotide comparison between the sequence of human RVA screening primers for VP7 (Beg9, sBeg9, 9Con1-L, EndA, VP7-Rdeg, End9 and RVG9) and VP4 (VP4-1-17F, Con3 and Con2) with their corresponding region of the new bat RVA genotype. Red nucleotide indicates dissimilar nucleotide between a strain segment sequence and primer and bold numbers at the beginning and end of sequence indicate nucleotide positions for different strains. For clarity the reverse complement sequence of the reverse primers is used for comparison with bat RVA sequences.

Overall, the similarity percentages for the VP7 forward and reverse primer between the bat RVA sequences and the human primers were 57.1-100% and 63.2-95%, respectively. For VP4, the percentage similarity ranged from 63.6-94.4% and 76.2-85.7% for the forward and reverse primers, respectively. Detail comparisons of each of the primer pair against each of VP4 and VP7 genotype described in this study is found in Suppl. S3.

Additionally, to determine if any of these bat RVAs could cross species and infect humans, we designed primers (RVA-VP6_40F and RVA-VP6_1063R) from an alignment of both human and bat RVA VP6 segments to screen 25 diarrheic infant samples (infants living around the same region where the bat samples were collected). Thirty-six percent of human samples were positive for RVA, however, none of them was of bat RVA origin. They all possessed the typical human genotype I1 and were 99% identical to the Gambian, Senegalese, Belgian and Brazilian Wa-like G1P[8] strains BE00007 (HQ392029), MRC-DPRU3174 (KJ752288), MRC-DPRU2130-09 (KJ751561), rj1808-98 (KM027132), respectively (Suppl. S4).

In summary, strain RVA/bat-wt/CMR/BatLy03/2014/G25P[43] possessed the genotype constellation G25-P[43]-I15-R16-C8-M15-A26-N8-T11-E23-H10 (Table 3), which shared six genotypes with those of the Kenyan bat RVA strain KE4852. For VP4 (P[6] vs P[43]), VP1 (R-unassigned vs R16) and NSP4 (E2 vs E23), different genotypes were observed, whereas for VP3 and NSP1 no sequence data were available for KE4852 for comparison. The 4 other strains were named BatLi10, BatLi08, BatLi09 and BatLy17 and possessed the genome constellations:

**Table 3:** Genotype constellations of all known bat RVAs. Pink indicates genotypes which are shared between Kenyan RVA strain KE4852 and BatLy03; red indicates (unknown) genotypes of KE4852; orange and blue indicate genotypes of Chinese bat RVA strains; and green shades represent all novel genotypes identified in Cameroonian bat RVAs.

| Strain | VP7 | VP4 | VP6 | VP1 | VP2 | VP3 | NSP1 | NSP2 | NSP3 | NSP4 | NSP5 |
| --- | --- | --- | --- | --- | --- | --- | --- | --- | --- | --- | --- |
| RVA/Bat-wt/KEN/KE4852/2007/G25P[6] | G25 | P[6] | I15 | Rx | C8 | Mx | Ax | N8 | T11 | E2 | H10 |
| RVA/Bat-tc/CHN/MSLH14/2012/G3P[3] | G3 | P[3] | I8 | R3 | C3 | M3 | A9 | N3 | T3 | E3 | H6 |
| RVA/Bat-tc/CHN/MYAS33/2013/G3P[10] | G3 | P[10] | I8 | R3 | C3 | M3 | A9 | N3 | T3 | E3 | H6 |
| RVA/Bat-wt/CMR/BatLi10/2014/G30P[42] | G30 | P[42] | I22 | R15 | C15 | M14 | A25 | N15 | T17 | E22 | H17 |
| RVA/Bat-wt/CMR/BatLi09/2014/G30P[42] | G30 | P[42] | I22 | R15 | C15 | M14 | A25 | N15 | T17 | E22 | H17 |
| RVA/Bat-wt/CMR/BatLi08/2014/G31P[42] | G31 | P[42] | I22 | R15 | C15 | M14 | A25 | N15 | T17 | E22 | H17 |
| RVA/Bat-wt/CMR/BatLy17/2014/G30P[xx] | G30 | P[xx] | I22 | R15 | C15 | M14 | A25 | N15 | T17 | E22 | H17 |
| RVA/Bat-wt/CMR/BatLy03/2014/G25P[43] | G25 | P[43] | I15 | R16 | C8 | M15 | A26 | N8 | T11 | E23 | H10 |

Gx-P[x]-I22-R15-C15-M14-A25-N15-T17-E22-H17 with G31P[42] for BatLi08, G30P[xx] for BatLy17, and G30P[42] for BatLi10 and BatLi09 (Table 3). Screening human samples for these bat RVAs indicated no interspecies transmissions and primer comparison showed that not all the strains can be picked up with the currently used screening primers.

## Discussion

Bats have been proven to harbor several human pathogenic viruses including SARS, MERS-related coronaviruses, as well as filoviruses, such as Marburgvirus, or Henipaviruses, such as Nipah and Hendra virus (13–15), but bat RVAs have only been sporadically reported. So far, only 3 bat RVA strains have been characterized and the first strain was reported in a straw-colored fruit bat in Kenya named RVA/Bat-wt/KEN/KE4852/2007/G25P[6] (17); while the other two, named RVA/Bat-tc/CHN/MSLH14/2012/G3P[3] and RVA/Bat-tc/CHN/MYAS33/2013/G3P[10], were isolated from a lesser horseshoe bat, and a Stoliczka’s trident bat in China, respectively (18, 19).

To better understand the spread and diversity of RVA in bats, we performed an RVA screening in Cameroonian bats, after trapping both male and female, young and adult bats close to human dwellings in Muyuka, Limbe and Lysoka localities of the South West region of Cameroon (Fig. 1). Using an unbiased viral metagenomics approach, we identified 5 divergent novel bat RVA strains, 4 of which were genetically similar to each other. The fifth strain was related to the Kenyan bat strain. Interestingly, all these RVAs were identified in adult (both female and male) straw-colored fruit bats (*Eidolon helvum*) which is in contrast to human and other animal whereby RVA (symptomatic) infections occur mostly in juveniles (1). Also, diarrhea or other obvious signs of sickness were not noticed in these bats. This may suggest that bats may undergo active virus replication and shedding without obvious clinical signs (33), which potentially could increase human exposure.

Even though there exists a considerable genetic divergence between bat RVA and human RVA, suggestions have been made about potential interspecies transmission of Chinese and Kenyan bat RVA strains. The two Chinese RVA strains are genetically quite conserved (all segments of both strains have the same genotype except for their VP4 gene). Based on genome comparisons of Chinese bat and partial human RVA strains from Thailand (CMH079 and CMH222) and India (69M, 57M and mcs60), Xia and colleagues speculated that Asian bat RVAs may have crossed the host species barrier to humans on a number of occasions (19). In addition, the unusual equine strain E3198 (34) shares the same genotype constellation with either MYAS33 and/or MSLH14 in all segments except VP6. This data therefore suggests that this equine RVA strain most likely share a common ancestor with Asian bat RVAs. Furthermore, the genotype constellations of these Asian bat RVA (Table 3) are reminiscent to the Au-1-like genotype backbone of feline/canine-like RVA strains, as well as to the genotype constellation of two unusual simian RVAs (RRV and TUCH) (35), suggesting that interspecies transmissions might have also occurred in the distant past. Moreover, an unusual Ecuadorean human RVA, Ecu534 (36) is closely related to bat sequences from Brazil recently submitted to GenBank. Similarly, possible interspecies transmission trends were also suggested by He and colleagues (18) between bovine strain RVA/Cow-wt/IND/RUBV3/2005/G3P[3]) and the bat strain MSLH14.

Given the novelty of the bat RVA strains described here, it is questionable if the currently used human RVA screening primers (for VP7 and VP4) will pick up these divergent strains in case an interspecies transmission from bats to humans would occur. Comparisons (Table 2 and Suppl. S2) of these primers with the corresponding sequences showed that the primer combination 9Con1-L and VP7-Rdeg (29, 30), would most likely detect both G25 and G31 RVA strains. Also, the combination of either Beg9 or sBeg9 with End9 or RVG9 (32) might be successful in amplifying G25, but the same combinations might not be able to pick up the novel G31 and G30 genotypes in PCR screening assays. Furthermore, the primer combination Beg9/sBeg9 and EndA (37) are likely to detect G25, but will be unsuccessful in case of G30 and G31 especially if the forward primer Beg9 is used. Considering VP4, both forward primers, VP4-1-17F (38) and Con3 (31) in combination with the reverse primer Con2 will be sub-optimal in detecting any of the P genotypes. Generally, with the exception of strain G25, amplification of most of the bat strains will be sub-optimally or not successful at all for the different available primer combination. Therefore, zoonotic events of bat RVA strains could easily be missed with the current screening primers depending on the primer combinations, PCR conditions and/or circulating zoonotic strains.

The genotype constellations of the two Chinese bat RVAs showed clear indication of recent reassortment event(s) because they possessed different P genotypes (P[10] for MYAS33 and P[3] for MSLH14), and some gene segments were nearly identical whereas others were not (18, 19). Their genotype constellation differs markedly from the Kenyan straw-colored fruit bat strain (KE4852). Although this strain showed a unique genotype backbone, some of its segments were similar to some human and other animal RVAs (17). Moreover, KE4852 share the same genotypes in several gene segments (VP2, VP6, VP7, NSP2, NSP3 and NSP5) with our bat RVA strain BatLy03 indicating possible reassortment events between different bat RVA strains, as well as a large geographical spread of this virus. Furthermore, BatLi08, BatLi09, BatLi10 and BatLy17 had conserved genotype constellations (in VP6, VP1-VP3, NSP1-NSP5) with 98-100% nucleotide sequence similarity except for the VP7 of BatLi08 and VP4 of BatLy17, again confirming reassortment events within bat RVAs.

In order to investigate the possibility of bat RVA infecting humans who are living in close contact with bats, we used novel primers (RVA-VP6_40F and RVA-VP6_1063R) designed from an alignment of both human and bat VP6 RVA segment to screen 25 infant samples from patients with gastroenteritis, living around the same region where the bat samples were collected. Interestingly, 36% of human samples were positive, however, none of these were positive for bat RVAs. All were of the typical human RVA genotype I1 and therefore there is no evidence for interspecies transmissions of bat RVA to humans. However, this result is not conclusive as only a small sample size was considered here. Sampling a larger number of subjects and from different localities around the region might result in more conclusive answers with respect to the zoonotic potential of these bat RVA strains.

The high genetic divergence and partial relatedness of most of the segments of the different bat RVA strains and the ones identified in this study indicate the frequent occurrence of reassortment events in the general bat population and those of Cameroon in particular. Also, with the current knowledge of the genetic diversity, there seems to exist several true bat RVA genotype constellations, as has been previously described for humans, and cats/dogs (10, 39). However, this needs to be further confirmed by identification of a larger number of RVAs from bats from different age groups and different geographical locations.

## Acknowledgements

KCY was supported by the Interfaculty Council for Development Cooperation (IRO) from the KU Leuven. NCN was supported by the Institute for the Promotion of Innovation through Science and Technology in Flanders (IWT Vlaanderen).

## Footnotes

**Competing interests** The authors declare that they have no competing interests.

**Authors’ contributions** KCY conceived and designed the study, collected samples, performed the experiments, analyzed the data and drafted the manuscript. MZ, NCN, EH, LB, WD and PM performed in the experiments, data analysis and contributed to manuscript drafting. SMG collected the samples and drafted the manuscript. JM and MVR conceived and designed the study and contributed to data analysis and manuscript drafting. All authors read and approved the final manuscript.

